# Correcting Preprocessing Bias in Sparse Chromatin Contact Data Enables Physically Interpretable Reconstruction of Genome Architecture

**DOI:** 10.64898/2026.03.31.715622

**Authors:** Stanislav Sys, Marcel Misak, Azza Soliman, Rosa Herrera-Rodriguez, Ruxandra-Andreea Lambuta, Stephan Weißbach, Michael Wand, Karin Everschor-Sitte, Susann Schweiger, Jasper J. Michels, Jan Padeken, Susanne Gerber

## Abstract

DNA is the largest biopolymer in nature, and chromatin contact maps are widely interpreted as quantitative readouts of its three-dimensional organization. However, the validity of such interpretations critically depends on how these maps are processed.

Here, we identify a previously overlooked but fundamental source of bias in chromatin contact data analysis. We demonstrate that a widely adopted preprocessing convention, namely whole-matrix percentile clipping, systematically distorts sparse contact maps by collapsing their dynamic range. This effect is strongest in near-diagonal interactions, precisely the regime encoding chromatin domains and looping structures, thereby compromising quantitative interpretation while preserving superficial structural features.

We show that this distortion represents a sparsity-dependent failure mode of current preprocessing standards and affects the comparability of datasets and computational methods across technologies and sequencing depths. To address this, we introduce a statistically consistent preprocessing framework based on nonzero-percentile clipping and log-space normalization, which preserves the intrinsic dynamic range of observed contacts. Building on this foundation, we present CCUT, a modular deep learning framework for chromatin contact map reconstruction. Under corrected preprocessing, reconstructed maps recover domain organization, contact decay, and scaling behavior consistent with polymer physics. Importantly, we demonstrate quantitative agreement between reconstructed maps and simulated contact patterns derived from a kinetic Monte Carlo loop extrusion model, enabling direct comparison between experimental data and physical models.

Together, our results establish preprocessing as a decisive determinant of the physical interpretability of chromatin contact maps and provide a principled framework for robust and comparable analysis across chromatin conformation capture technologies.

## Introduction

Chromatin contact maps are widely interpreted as quantitative readouts of genome organization, yet their physical interpretability critically depends on statistical preprocessing choices that are rarely examined. Despite their interpretation as physical measurements of an underlying polymer system, the statistical processing of these data has not been systematically aligned with this interpretation.

Conformation Capture techniques (3C) such as Hi-C have become fundamental tools for investigating higher-order genome organization, enabling the mapping of structures that influence gene expression through their three-dimensional configuration in the nucleus (Lieberman-Aiden et al., 2009; Rao et al., 2014; Rao et al., 2017; Zhong et al., 2023). Such methods measure the interaction frequencies between different loci in the genome and can be used to locate specific structures such as A/B compartments, chromatin loops, or topologically associated domains (TADs) in the genome. These secondary DNA structure features can influence gene expression via their spatial configuration within the nucleus (Zhong et al., 2023). More recent methods such as Micro-C (Krietenstein et al., 2020) allow for a more fine-grained picture of the spatial organization of chromatin in the nucleus. The reconstruction of the three-dimensional arrangement of these components is essential for deciphering the relationships between chromatin organization and genetic function (Krietenstein et al., 2020; Lieberman-Aiden et al., 2009; Rao et al., 2017; Wang et al., 2022; Zhong et al., 2023). Despite the advances achieved using these approaches, 3C-based methods still face obstacles in the characterization of higher-order chromatin structures. These limitations largely derive from canonical short-read sequencing biases: nonuniform coverage across GC-rich or GC-poor loci, diminished detection of large structural rearrangements, and ambiguous read alignment within repetitive DNA (Browne et al., 2020; Roberts et al., 2021; Weissbach et al., 2021). These challenges are discussed in detail in excellent reviews by (Li & Durbin, 2024; Zhang et al., 2024). Pore-C aims to close this gap and has been used extensively to generate new insights by leveraging Oxford Nanopore long-read sequencing technology (Chemparathy et al., 2024; Deshpande et al., 2022; Roberts et al., 2021; Zhong et al., 2023). It provides an additional layer of information compared to classical 3C methods by providing access to global high-order multiway contacts and capturing methylation information at the same time (Deshpande et al., 2022). These advancements are fundamental, as gene regulation often entails subtle chromatin interactions across multiple loci (Handoko et al., 2011; Zhong et al., 2023), which at least in part may occur in selective nuclear condensates (Hnisz et al., 2017; Sabari et al., 2018). Together Pore-C has the chance to provide a detailed characterization of higher order nuclear organization, overcoming the limitation of being restricted to pairwise contacts and providing a higher resolution map of repetitive, heterochromatic regions. Its major drawback however is the cost of nanopore sequencing associated with the need for deep coverage in 3C based experiments.

Current computational enhancement methods implicitly inherit preprocessing conventions that were originally established for dense Hi-C datasets, without re-evaluation for sparse data regimes, namely a preprocessing pipeline originating in DeepHiC (Hong et al., 2020): Knight-Ruiz-balancing followed by clipping at a fixed threshold of 255, a value corresponding to the approximate 99.9th percentile of KR-balanced 10 kb contact counts that is specific to this specific dataset generated in the GM12878 cell line (Rao et al., 2014). This threshold was adopted by HiCARN (Hicks & Oluwadare, 2022), HiCSR (Dimmick et al., 2020), and ScHiCAtt (Menon et al., 2025). HiCDiff (Wang & Cheng, 2024) and SHICEDO (Huang et al., 2025) differ in their assumptions: HiCDiff uses percentile clipping on ICE balanced matrices and SHICEDO adopts the distance stratified BandNorm, that is specific to single cell 3C experiments. While this convention might be reasonable for Hi-C, where count distributions are relatively stable across experiments, it is not statistically appropriate as a general strategy. As we demonstrate below, 3C technologies differ dramatically in their count distributions: Pore-C contact matrices are substantially sparser in comparison to Hi-C or Micro-C contact matrices, with a higher proportion of zero entries and their non-zero count distributions are highly sensitive to sequencing depth.

Applying whole-matrix percentile clipping under these conditions creates a shifted distribution with high zero counts and thus shifts the clipping thresholds to a relatively small value. This represents a sparsity-dependent failure mode, in which the dominance of zero entries shifts percentile thresholds into the bulk of the observed signal. The clipping value scales with the sparsity of the matrix, which in turn depends on the sequencing technology and depth. This means that a preprocessing protocol optimized for one experiment type will systematically distort data from another, and that performance comparisons between methods are confounded whenever preprocessing choices differ.

Compressing contact counts into a narrow range is a practical requirement for min-max scaling to [0, 1], which empirically stabilizes neural network training. The clipping can therefore serve as a workaround for high dynamic ranges and heavy-tailed distributions of raw contact data. Nonetheless, this workaround comes at the cost of discarding a big proportion of the signal structure and it makes a lossless or near-lossless reversal to original count space essentially impossible. We resolve this by introducing a log-space normalization pipeline that uses nonzero-percentile clipping at p99.95, followed by a log1p transform, after which per-chromosome min-max scaling to [0, 1] is applied using per-chromosome percentiles and maxima. The resulting predictions are thus directly interpretable in terms of the original contact counts.

Here, we demonstrate that preprocessing is a decisive determinant of chromatin contact map fidelity and introduce a framework that restores their physical interpretability. First, we characterize how whole-matrix clipping severely degrades pixelwise similarity metrics in matched comparisons and is specifically harmful to the near-diagonal signal by flattening the near diagonal values, representing close range interactions. Nonzero-percentile clipping, which restricts the percentile calculation to observed contacts and thereby preserves the natural dynamic range of the data, eliminates this distortion and yields results consistent with raw unclipped matrices. Second, we introduce the Chromatin Capture Upsampling Toolbox (CCUT), a modular deep learning framework centered on HiCNet, a hierarchical network architecture adapted for chromatin contact matrix enhancement. CCUT is designed to be trained and evaluated under statistically appropriate preprocessing. We demonstrate its application to Pore-C as a primary use case, as no dedicated computational enhancement tool currently exists for this technology. We show that HiCNet faithfully recovers contact decay, insulation structure, and TAD boundaries from deeply downsampled Pore-C data, and we show that Lin’s concordance correlation coefficient (CCC) provides a more informative measure of contact decay reconstruction than Pearson correlation, which saturates near unity even for substantially degraded matrices due to the monotonic nature of P(s) curves. Given how especially close range interactions are dictated by the behavior of chromatin as a polymer we show that the upsampled interaction maps are consistent with simulated matrices obtained using a loop extrusion kinetic Monte Carlo (KMC) model that combines polymer physics with dynamic association of extrusion motors and boundary proteins. Illustrating the universal adaptability of our approach, we applied CCUT to Pore-C interaction maps of the around ∼300x smaller, but highly dense *C. elegans* genome.

## Results

### Whole-matrix clipping collapses the dynamic range of sparse contact data and selectively distorts near-diagonal signal

To illustrate that 3C methods are not comparable in their raw count distributions, we first examined chr1 at 50 kb resolution across representative Hi-C, Micro-C, and Pore-C datasets (Figure 1a). In the representative datasets we picked for this analysis the fraction of zero entries increases progressively from 33.1% in Hi-C to 50.8% in Micro-C to 94.6% in Pore-C, reflecting the fundamental difference in sequencing depth and ligation efficiency between these technologies. As a direct consequence, the whole-matrix p99.95 clipping threshold, which includes zero counts in the percentile calculation, collapses toward the lower end of the observed count distribution as sparsity increases. For Pore-C, this threshold reaches 24 counts, 45-fold below the nonzero p99.95 threshold of 1070. For reference, several widely used enhancement tools apply either whole-matrix clipping at p99.9 like HiCDiff, placing their thresholds similarly deep within the bulk of the Pore-C count distribution, while others use a hardcoded cutoff of 255 as HiCARN (Figure 1a). Nonzero-percentile clipping on the other hand restricts the percentile calculation to observed contacts, preserving the natural dynamic range of the data without the heavy impact of zero values. Thus, nonzero-percentile-clipped data is robust to differences in sparsity across technologies and sequencing depths.

**Figure 1:**
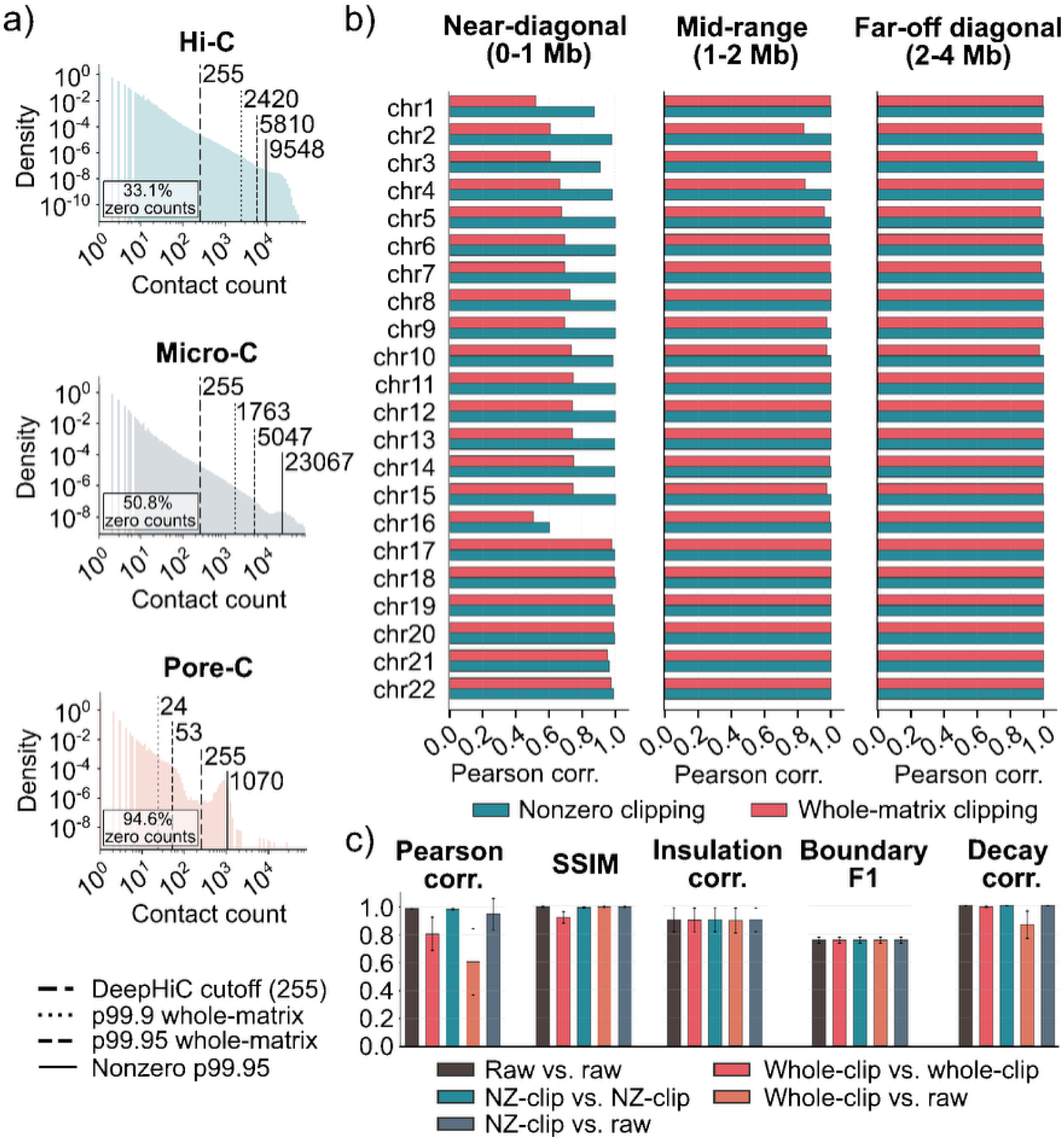
Whole-matrix clipping collapses the dynamic range of sparse contact data and selectively distorts near-diagonal signal. **(a)** Contact count distributions (nonzero entries, log-scale density) for chr1 at 50 kb resolution across three chromatin capture technologies.

Vertical lines indicate clipping thresholds in absolute count values: nonzero p99.95 (solid), whole-matrix p99.95 (dashed), p99.9 whole-matrix as used by several existing tools (dotted), and the DeepHiC’s hardcoded cutoff of 255 (long dash). Zero fractions are annotated per panel. **(b)** Per-chromosome Pearson correlation coefficients clipped (p99.95) and non-clipped contact matrices, stratified by genomic distance into near-diagonal (0-1 Mb), mid-range (1-2 Mb), and far-off-diagonal (2-4 Mb) regions. Teal bars indicate nonzero-percentile clipping, and coral bars indicate whole-matrix clipping. **(c)** Comparison of five evaluation metrics: Pearson correlation coefficient of the flattened contact matrix (PCC), structural similarity index measure (SSIM), insulation score correlation (PCC of insualtion score), topologically associating domain (TAD) boundary detection F1 score, and Pearson Correlation of the contact decay correlation. Calculated between 16x binomially downsampled and high-resolution contact matrices. The analysis contrasts three preprocessing pipelines: raw (unclipped), whole-matrix clipping at the p99.95 percentile, and nonzero clipping at the p99.95 percentile. Bars represent mean values across 22 autosomes with standard deviation error bars.

To quantify the consequences of this dynamic range collapse for contact matrix fidelity, we computed the Pearson correlation coefficient between whole-matrix clipped and nonzero-clipped matrices against their unclipped originals, stratified by genomic distance band across all 22 autosomes (Figure 1b). The information loss from whole-matrix clipping is strongly concentrated in the near-diagonal band (0-1 Mb), where high-count contacts between proximal loci reside. For several chromosomes, near-diagonal Pearson Correlation Coefficient (PCC) falls to 0.5-0.7 under whole-matrix clipping, while nonzero clipping maintains PCC above 0.85 genome-wide. In the mid-range (1-2 Mb) and far off-diagonal (2-4 Mb) bands, both clipping methods perform similarly, with PCC values approaching 1.0 for most chromosomes. This distance dependence is expected: whole-matrix clipping specifically truncates the high-count near-diagonal entries that dominate the sparse tail of the count distribution, leaving low-count long-range contacts largely unaffected. This is precisely the region that encodes biologically meaningful contact domains such as TADs and chromatin loops, whose distortion under whole-matrix clipping directly compromises the signal that enhancement models are trained to recover.

We next evaluated the downstream effect of clipping methods on a comprehensive panel of similarity metrics, comparing 16× binomially downsampled matrices against their full-coverage counterparts under three preprocessing schemes: raw, whole-matrix p99.95 clipping, and nonzero p99.95 clipping (Figure 1c). Whole-matrix clipping produces a degradation of pixelwise metrics in matched comparisons: mean PCC across 22 autosomes drops from 0.986 (raw vs. raw) to 0.807 (whole-clipped vs. whole-clipped), and Structural Similarity Index (SSIM) drops from 0.993 to 0.918. Nonzero clipping recovers near-raw fidelity, with PCC of 0.984 and SSIM of 0.989. In contrast, higher-order structural metrics are robust in terms of clipping methods: insulation score correlation remains at 0.900, TAD boundary F1 at 0.755-0.756, and contact decay correlation at 1.000 across all three matched preprocessing conditions. This dissociation has a practical implication in both directions: biological conclusions drawn from whole-clipped data regarding domain boundaries and compartment structure are likely reliable, but quantitative enhancement benchmarks reported under whole-matrix clipping significantly underestimate true model performance and are not comparable across studies that differ in preprocessing choice. We also note an apparent anomaly in the SSIM results: the cross-preprocessing comparison of whole-clipped downsampled versus raw high-resolution matrices yields SSIM of 0.993, higher than the matched whole-clipped comparison (0.918). This is a known artefact of SSIM’s sensitivity to dynamic range (Dosselmann & Yang, 2011). The stability constants in the SSIM formula scale with the observed data range L; whole-matrix clipping compresses L severely, making these constants small and magnifying the relative contribution of any residual difference between matrices. This behavior illustrates why cross-preprocessing SSIM comparisons are unreliable and further complementary metrics should be considered.

### CCUT recovers chromatin structure from deeply downsampled Pore-C data

Having established the importance of statistically appropriate preprocessing, we trained HiCNet within the CCUT framework on publicly available Pore-C data (Deshpande et al., 2022) and evaluated reconstruction quality on an independently produced in-house HG002 Pore-C dataset, processed using the nonzero log-space pipeline. We assessed reconstruction quality on two representative genomic regions on chr8 and chr20: comparing original high-resolution matrices against 16x downsampled and HiCNet-enhanced outputs (Figure 2). HiCNet recovers the characteristic triangular domain structure and off-diagonal interaction patterns that are almost entirely absent in the downsampled matrices, where contact signal is dominated by the main diagonal. This recovery is consistent across both regions and is reflected quantitatively across pixelwise, structural, and decay metrics.

**Figure 2:**
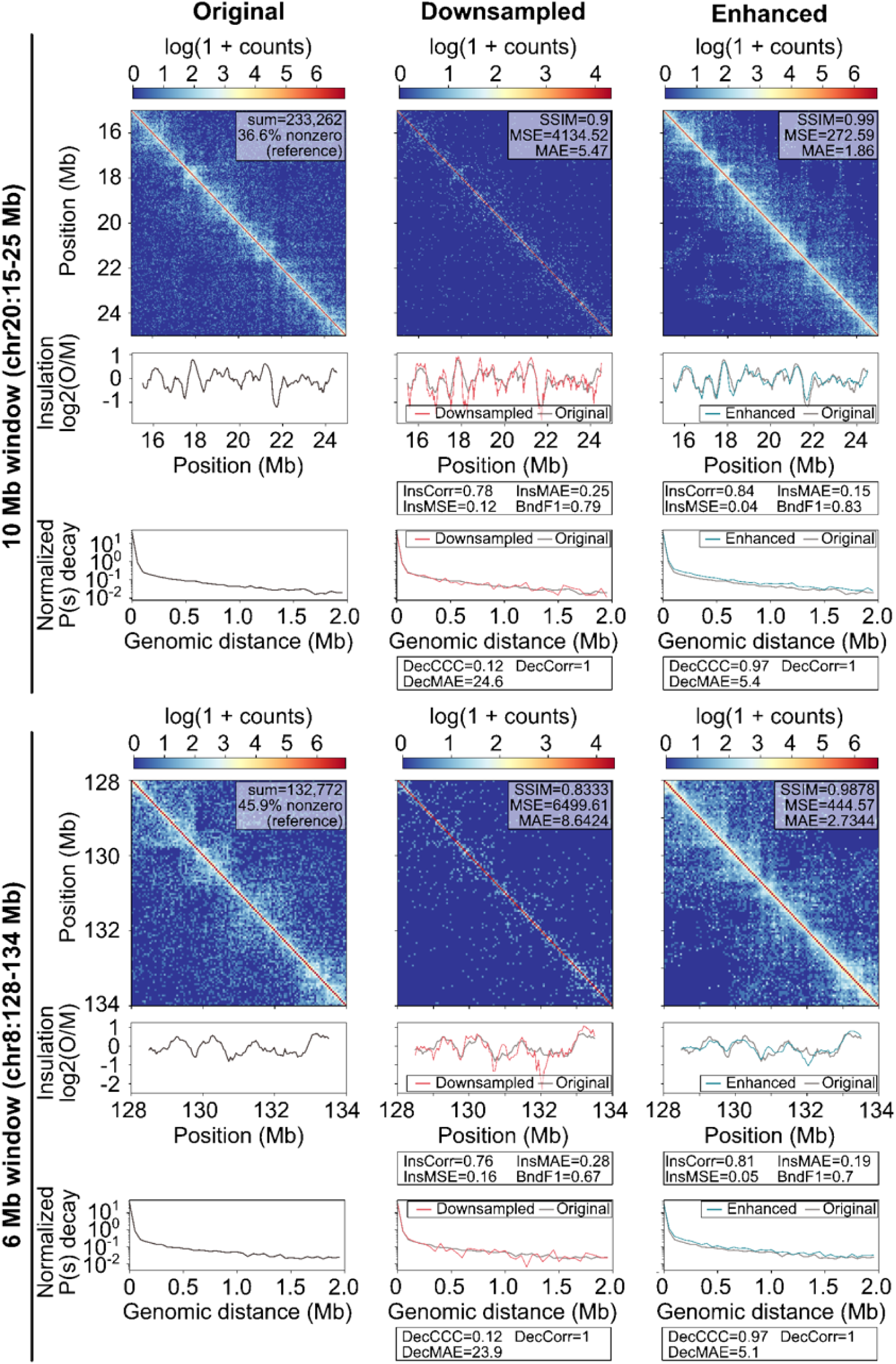
Visual and quantitative assessment of contact matrix reconstruction on representative genomic loci from the HG002 dataset. Each block shows three columns (original high-resolution, 16x binomially downsampled, and HiCNet-enhanced) and three rows (contact heatmap, insulation score profile, and contact decay curve). Top block: chr20:15-25 Mb (10 Mb window, 36.6% nonzero entries, sum=233,262). Bottom block: chr8:128-134 Mb (6 Mb window, 45.9% nonzero entries, sum=132,772). Bottom Contact matrices are shown as log(1+count) and were preprocessed using nonzero p99.95 clipping followed by log1p normalization. Pixelwise metrics (SSIM, MSE, MAE) are annotated on each heatmap relative to the high-resolution reference. Insulation score profiles (log2 observed/mean) are shown with insulation correlation, MAE, MSE, and TAD boundary F1 score annotated. Contact decay curves P(s) are shown on a log scale normalized to the genome-wide mean, with Lin’s concordance correlation coefficient (CCC), Pearson correlation, and MAE annotated.

For the subregion of chr8, SSIM improves from 0.833 in the downsampled matrix to 0.988 in the enhanced output, with MSE dropping from 6499.61 to 444.57 and MAE from 8.64 to 2.73. Insulation score correlation increases from 0.759 to 0.805, insulation MAE decreases from 0.280 to 0.187, and TAD boundary F1 improves from 0.667 to 0.700. For chr20, SSIM improves from 0.898 to 0.993, MSE from 4134.52 to 272.59, and MAE from 5.47 to 1.86, with insulation correlation increasing from 0.780 to 0.844 and boundary F1 from 0.791 to 0.833. The consistent improvement across two chromosomes on different genomic scales, differing gene density (chr20: 4.5 genes/Mb vs. chr 8: 8.7 genes/Mb), and domain organization suggests that HiCNet generalizes across genomic contexts rather than overfitting to specific regional features.

Recovery of the contact decay profile P(s) is a particularly stringent test of enhancement quality because it is sensitive to both the shape and the absolute level of the decay curve. Pearson correlation alone is insufficient for this assessment: due to the monotonic nature of P(s) decay, both the downsampled and enhanced matrices yield Pearson correlations near 1.0 with the original. Lin’s concordance correlation coefficient (CCC), which jointly measures correlation and agreement in scale and location, discriminates effectively between the two. For chr8, CCC for the downsampled matrix is 0.12, reflecting the severe amplitude mismatch between the sparse downsampled decay curve and the original, while CCC for the enhanced matrix reaches 0.97. This pattern replicates on chr20, with downsampled CCC of 0.12 and enhanced CCC of 0.97. We therefore recommend CCC as the primary metric for evaluating P(s) reconstruction in chromatin enhancement studies.

### Outlook and application: comparison with simulated multi-cell contact maps

The availability of CCUT as a tool capable of accurately reconstructing domain boundaries and enabling reliable TAD calling allows for developing high-level mechanistic understanding of chromatin (re)organization at the level of the dynamics of cohesin and boundary proteins, such as CTCF. To provide a proof-of-concept, we develop a 1D chromatin extrusion model in the spirit of the seminal work by Alipour and Marko (2012) and Goloborodko et al. (2016). The model should produce multi-cell contact matrices, which we aim to directly compare to upsampled experimental maps. For this, we explicitly allow for reversible binding of extrusion motors (“cohesin”) and boundary proteins (“CTCF”), characterized by appropriate rate constants, i.e. 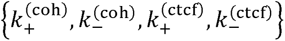. The cohesins are sampled from a fixed size reservoir, whereas the CTCF dynamics is captured through the occupancy of pre-fixed but randomly assigned binding sites with a given orientation. In steady state the fractional binding/occupancy is given by: 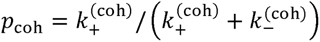 and 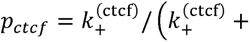 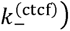.

Once a cohesin complex, which, for simplicity we only treat in a bound and unbound state, binds to the chromatin, extrusion may occur based on the following drift-diffusion formalism:

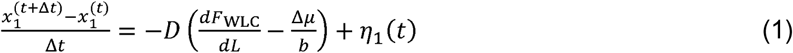

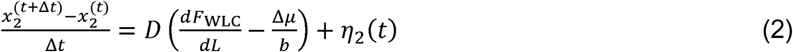

with *x*_1_ and *x*_2_ the genomic positions of the two heads of the cohesin complex on a strand of chromatin comprising *N* bins. *D* is a diffusivity and *L(t)* the size of the extruded loop at time *t*. A model with coupled head dynamics, as represented by Equations 1 and 2, is less advanced than a model with independent extrusion, but, as we will demonstrate below, for the purpose of this work it suffices to reproduces prominent features of the experimental (upsampled) contact matrices. *F_WLC_* is the looping free energy of the chromatin given according to the worm-like chain (WLC) model, as a linear combination of contributions due to translational entropy and bending stiffness, interpolated by an approximate cross-over correction accounting for semiflexibility (Giovan et al., 2015):

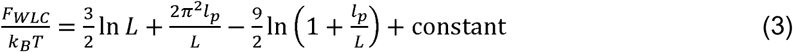

with *l_p_* the persistence length. The force 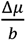 represents an ATP-driven extrusion bias, with a fundamental lengthscale. All lengthscales are non-dimensionalized by normalizing them by the chromatin length associated to one bin. In effect, this implies that the scale at which the model operates is determined by the value of the persistence length. Finally in Equations 1 and 2, *η(t)* represents thermal noise, with amplitude given by the fluctuation-dissipation theorem: <*η(t)η(t’)*>=*2Dδ(t*-*t’)* During time stepping, the strength of a contact increases as long as it is supported by a loop, while considering i) the dependence of the looping probability on the distance *s=j - i* between two genomic loci *j≥ i* and ii) the translational invariance of the loop geometry. The latter consideration is essential to the appearance of corner peaks in the calculated contact maps once both cohesin extrusion heads stall at convergently oriented CTCF sites. We define the looping statistical weight as:

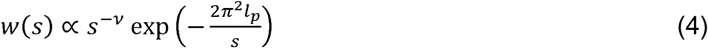

where the scaling exponent *v* may vary depending on the chromatin properties and the considered length scale. For instance, *v* = 3/2 for an equilibrium Gaussian chain but usually expresses flattened behavior (*v* < 3/2) during loop extrusion (Fudenberg et al., 2016). The exponential factor becomes important if the separation between the loci is so small that mutual contact is prohibited by the bending stiffness of the chromatin.

After updating the positions of the heads of a cohesin, the contact strengths within the corresponding loop are hence updated as:

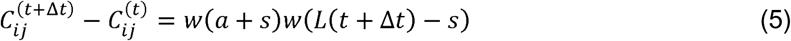

with α an off-set which avoids division by zero if the genomic loci fall into the same bin. If its value is small (≤1), deviation from the power law in the distance dependence of the contact probability remains limited to very small loops. To mimic the multi-cell (cell-averaged) nature of the experimental contact maps, our model groups and averages instantaneous snapshots, taken at regular intervals, while normalizing the contact strengths by the number of time steps. Our model allows for an exponential decrease in contact strength, consistent with diffusive decay of a loop upon cohesin dissociation. During simulation we keep track of how many loops support a given contact. The contact does not decay prior to the release of the last loop it is supported by.

Next, we demonstrate how the KMC model reproduces features observed in experimental (upsampled) data. For this, we consider the genomic loci from the HG002 data set depicted in Figure 2. These maps reveal various stripe patterns consistent with the occurrence of TAD boundaries, where extrusion by cohesin stalls at CTCF sites. The challenge now is to reproduce these relatively sharp features in the simulations despite the time averaging and taking into account the much more dynamic nature of boundary proteins compared to cohesin. In the simulations we adopt the same genomic extent (6 Mb) and resolution as for chr8:128-134 Mb binned at 50 kb (Figure 2, top left panel), i.e. producing a 120×120 contact matrix. For all calculations, we fix the mean density of CTCF binding sites at one site per ∼133 kb and ensure a steady state occupancy of one cohesin per ∼136 kb, based on a resolution of 50 kb per bin and consistent with reported densities (Fudenberg et al., 2016; Oti et al., 2016). Table 1 lists all input parameters used in the simulations.

**Table 1:**
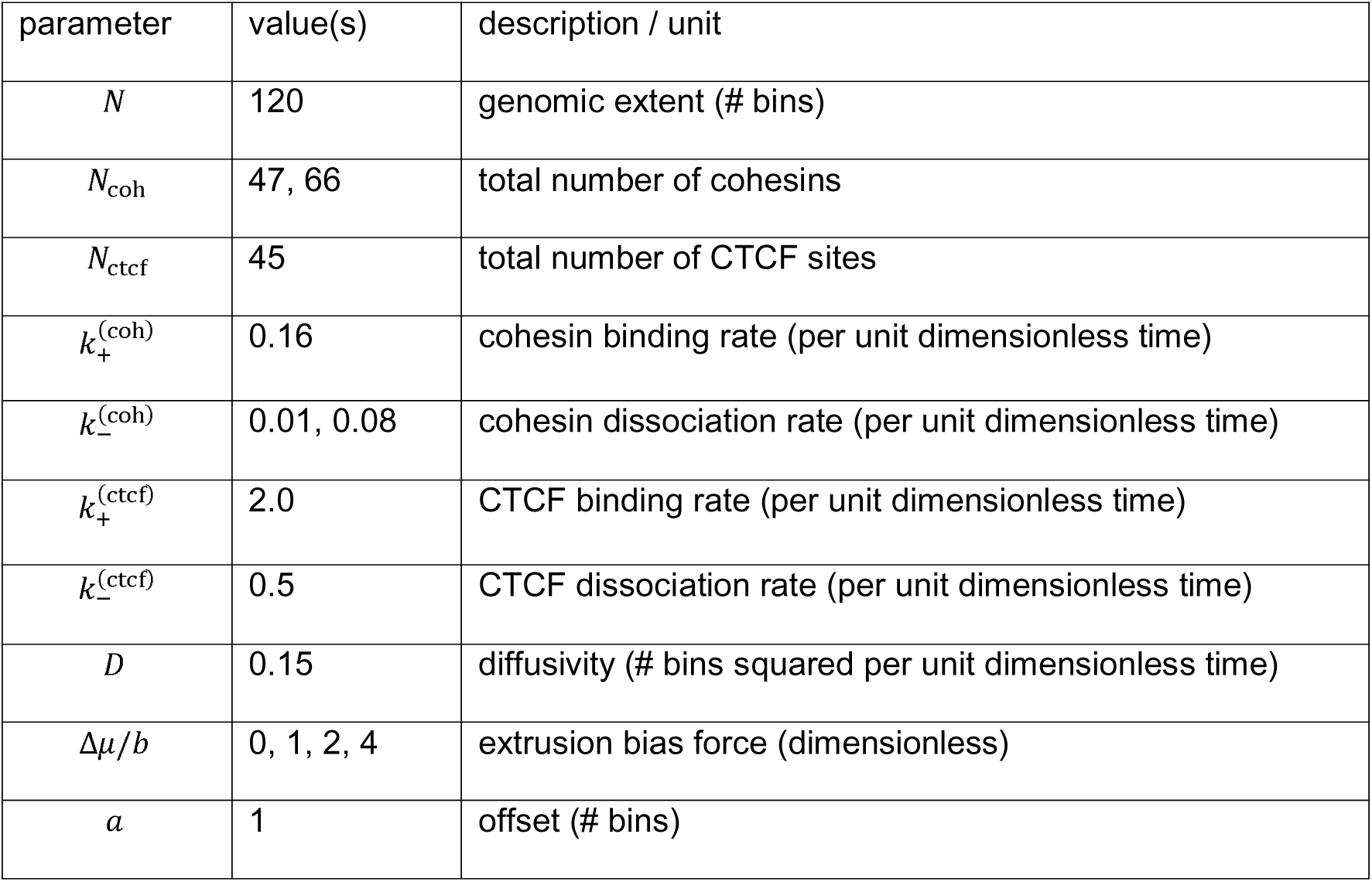
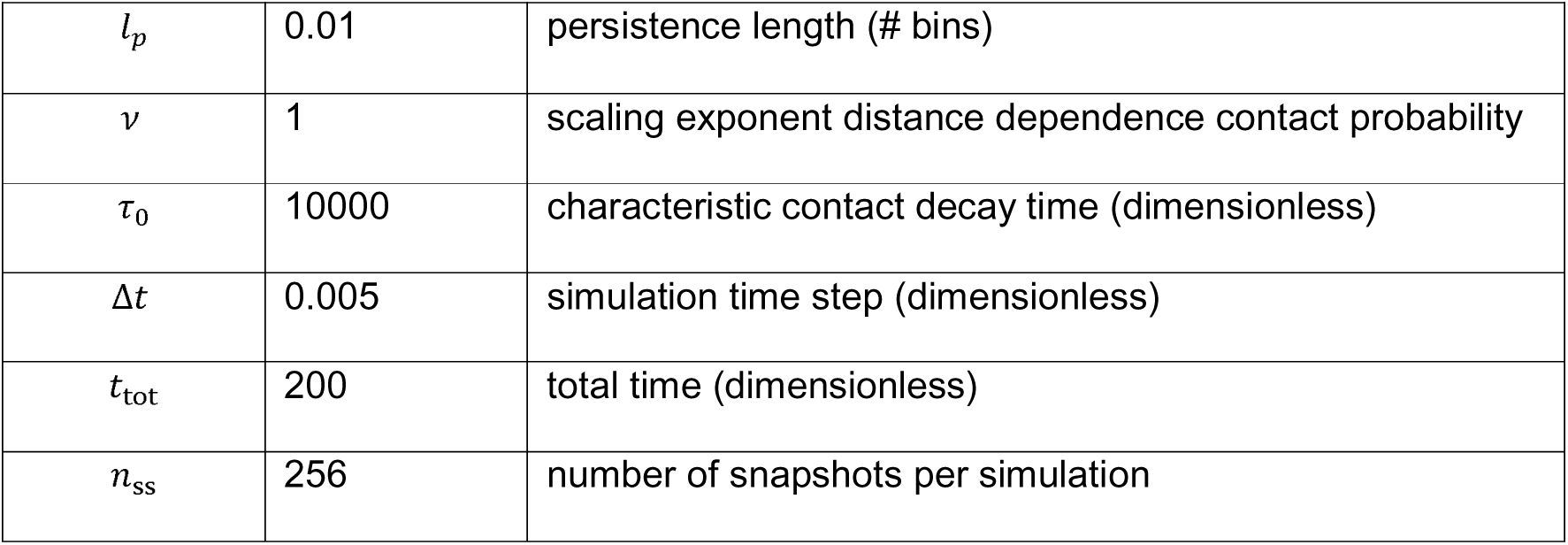
Input parameters for the KMC loop extrusion model.

We perform the simulations for increasing extrusion bias force *Δµ/b* and increasing cohesin dissociation rate constant 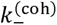. In biology, variation in these parameters may for instance be caused by fluctuation in the local concentration of ATP and/or cofactors that modulate cohesin unloading, such as WAPL (wings apart-like protein) (Gandhi et al., 2006). Figure 3a shows that the fractional occupancy of both CTCF and cohesin reaches steady state fast on the time scale of the simulations, irrespective of 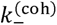. Naturally, for a high 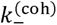 (Figure 3c), the steady state fractional occupancy is suppressed, but this is compensated for by a larger total number of available cohesins (Table 1).

**Figure 3:**
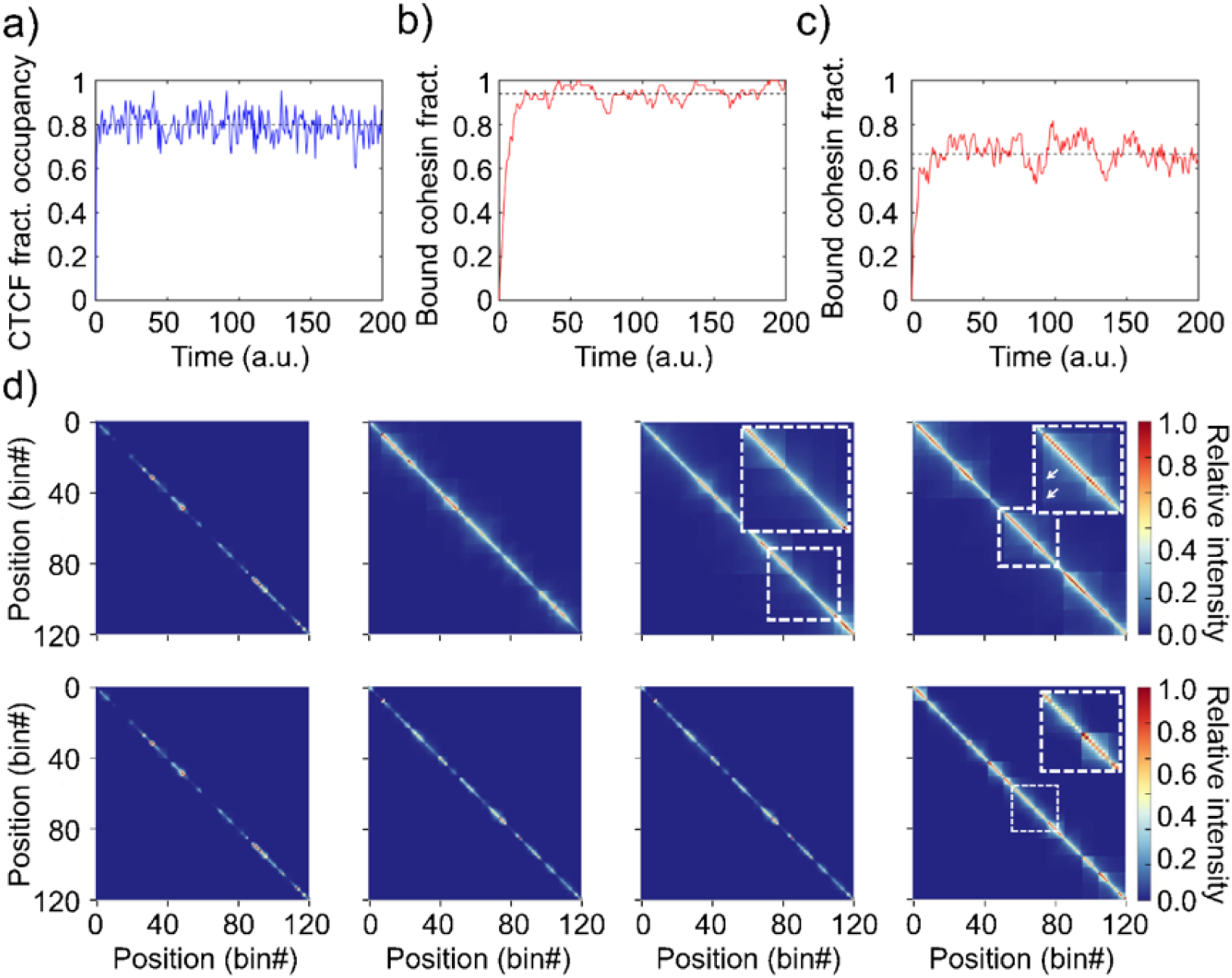
Simulated contact maps with TAD-related features. **(a)** Fraction of occupied CTCF binding sites as a function of (dimensionless) simulation time. **(b,c)** Fraction of bound cohesins for, respectively, slow and fast dissociation (see **Table 1**). The horizontal dashed black lines indicate the steady state values. **(d)** Representative steady state snapshots of simulated time-averaged contact matrices, for (left to right) an increasing extrusion bias (see **Table 1**) for slow (upper row) and fast (bottom row) cohesin dissociation. The insets show magnified regions as indicated by the dashed boxes to respectively highlight i) ‘chevron’-like features arising from stalling at CTCF sites with similar orientation, ii) enhanced corner peaks and iii) cross-like stripes due to stalling at neighboring boundary sites with opposite orientation. The color scale linearly interpolates between zero and the highest value in each matrix.

The simulated contact matrices (Figure 3d) show that an increase in extrusion bias leads to larger contact domains owing to more extensive directional extrusion, which is further enhanced if cohesins have a longer residence time due to a lower dissociation rate. Most maps clearly show stripes resulting from one-side stalling of extrusion at boundary proteins. These stripes, which are also seen in the experimental contact maps in Figure 2, increase in length with increasing bias and are most pronounced in the right most maps of the top row. The insets highlight regions containing ‘chevron-like’ and cross-like stripe patterns, which both can be discerned in the experimental maps in Figure 2. The inset in rightmost map of the top row shows enhanced corner peaks resulting from two-sided stalling at convergently oriented boundary proteins (Ghosh & Jost, 2020).

To test the versatility of CCUT we reconstructed the interaction map of *C. elegans* embryos. With 100Mb the *C. elegans* genome is ∼300x smaller than the human genome, but encodes for a similar number of genes (19,984 coding genes, WS290). The resulting gene density of ∼5-6 genes per 10 kb provides a challenge to any 3C based method and provides a real-world application for CCUT. Extensive chromatin profiling has shown that the holocentric *C. elegans* autosomes can be, at low resolution, subdivided into two heterochromatic arms, enriched for H3K9me, that flank a central domain enriched for actively expressed genes (Ahringer & Gasser, 2018). The interaction maps reconstructed by CCUT can clearly recapitulate this broad compartment organization, as exemplified by the reconstruction of chromosome 3 (Figure 4a). It also allows the detection of smaller TAD-like structures at the transition of eu- and heterochromatin. A second well described feature of the *C. elegans* genome is the separation of the X chromosome into smaller TAD-like domains by the dosage compensation complex (DCC) (Anderson et al., 2019; Das et al., 2024; Meyer, 2018). Similar to the bigger compartments does reconstruction of the X chromosome by CCUT allow for the identification of these TAD-like domains from sparse Pore-C data (Figure 4b).

**Figure 4:**
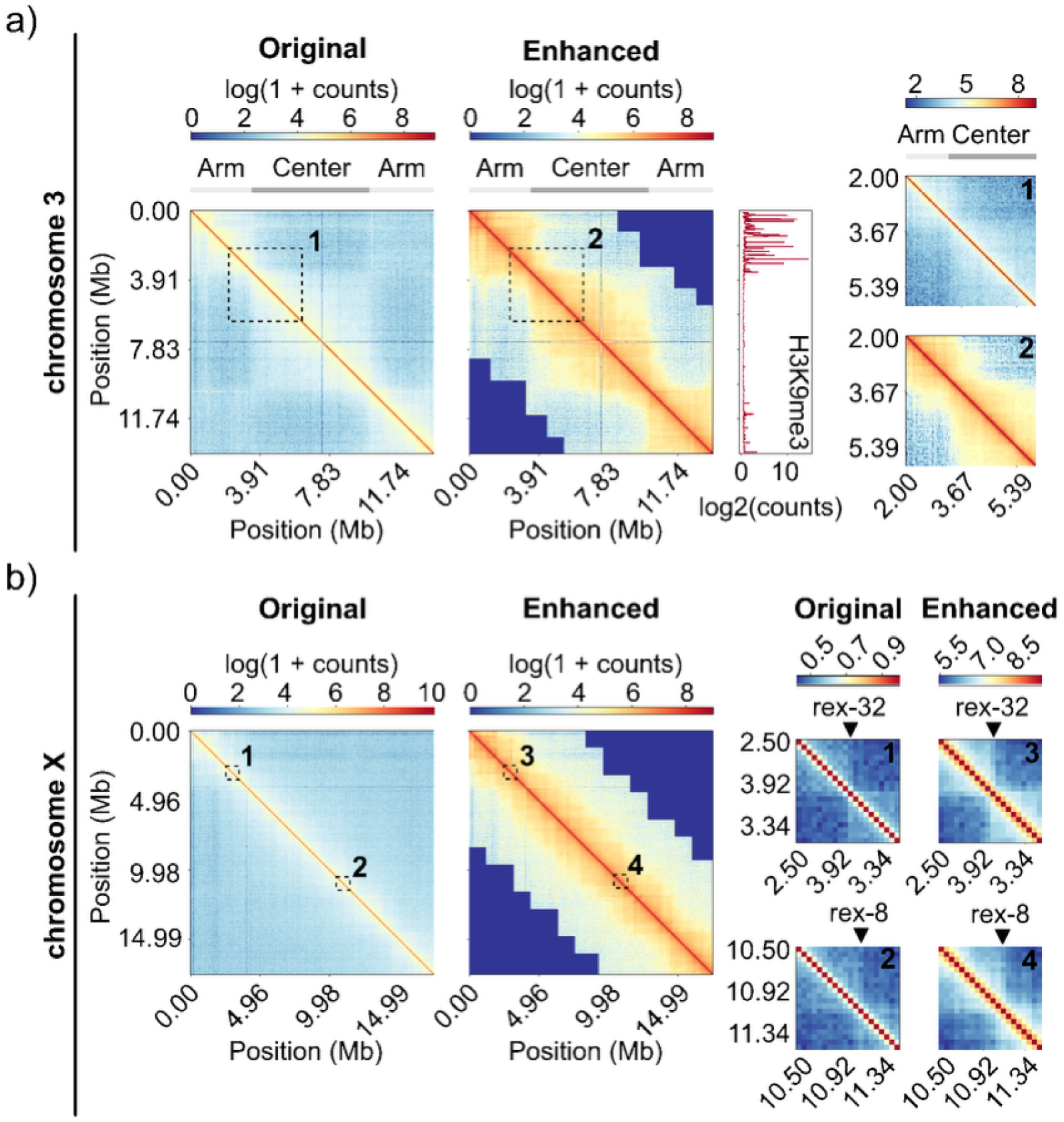
Comparison of original and reconstructed interaction maps of the highly dense *C. elegans* genome. **(a)** Original and reconstructed interaction map of chromosome III with the typical separation into heterochromatin rich chromosome arms (light grey) and euchromatin rich center region (dark grey) indicated below the heatmaps. Enrichment of heterochromatin marked by H3K9me3 for chromosome III measured by CUT&Tag in wild-type *C. elegans* embryos is plotted right to the enhanced interaction map. The right side shows a comparison of interaction maps of the eu-/heterochromatin border on the left arm on chromosome 3 for original and reconstructed data. **(b)** Original and reconstructed interaction map of chromosome X with the typical separation into smaller TAD-like domains organized by the C. elegans dosage compensation complex (DCC). Enlargements showing the reconstructed interaction frequencies at TAD boundaries created by experimentally validated recruitment sites of the DCC (rex-32, rex-8).

The above exercise is just to demonstrate the pipeline we have in place to extract mechanistic conclusions via upsampling of low coverage data. Although an in-depth mechanistic study including variation in experimental conditions is out of the scope of this work, we emphasize that the KMC model is readily extended with additional processes, for instance including reaction networks involving cofactors that regulate chromatin extrusion, as well as cohesin and CTCF binding and dissociation. It is furthermore noted that although a model is per definition based on simplifying assumptions, it is either inherently noise-free or amenable to accommodating controlled levels of noise. Hence, simulations may in principle also be used to help discriminate between biophysically meaningful contact patterns and noise-related features.

## Discussion

Our results demonstrate that preprocessing is not a technical detail, but a determinant of whether chromatin contact data can be interpreted as physically meaningful measurements. Computational enhancement methods offer a practical route to reducing these costs, but their development has been shaped by preprocessing conventions that were established for specific experimental contexts. To our knowledge, these methods have not been systematically re-evaluated as the field has expanded to sparser data types. Here, we address this gap by systematically evaluating how preprocessing choices affect contact matrix fidelity and enhancement benchmarking, and by introducing CCUT, whose predictions are consistent with polymer models of chromatin organization.

### Preprocessing conventions are technology-specific

The preprocessing pipeline that has dominated the field since its introduction normalizes contact matrices by clipping at a fixed percentile or hardcoded maximum, followed by min-max scaling to the unit interval. This convention was developed and validated on dense Hi-C datasets where the fraction of zero entries is moderate and the whole-matrix count distribution is reasonably well-behaved compared to Pore-C. Under these conditions, whole-matrix percentile clipping produces a threshold that falls sensibly within the tail of the nonzero count distribution, and the resulting distortion of the dynamic range is limited. The strong performance reported by existing methods on these datasets reflects both genuine model quality and the fact that the preprocessing artefacts we characterize here are relatively mild when sparsity is low.

We observe that with increasing matrix sparsity zero entries start to dominate the count distribution, as they do in Pore-C data where up to 94.6% of bins are unobserved at typical sequencing depths. The whole-matrix percentile thresholds impact is highly influenced by the number of zero counts and collapses into the bulk of the nonzero signal. As we demonstrate, this produces clipping thresholds that fall orders of magnitude below those obtained by restricting the percentile calculation to observed contacts for all three sequencing paradigms, especially for Pore-C. The consequence is not simply a cosmetic change to the matrix appearance but that the near-diagonal high-count entries that encode TAD structure and chromatin loops are truncated to near-zero. A model trained to map a clipped-to-noise low resolution matrices onto equally clipped high-resolution targets is bound to learn a noisy chromatin structure representation.

This sparsity-dependence explains why the problem went unnoticed in the Hi-C enhancement literature. On the datasets where these methods were developed and benchmarked, the distortion of metrics like SSIM and insulation scores is small enough that models still learn useful mappings and metrics still reflect genuine improvement. Importantly, this effect is not specific to Pore-C, but reflects a general sparsity-dependent limitation of percentile-based normalization strategies. It is only when the same conventions are applied to substantially sparser data as sparse, deeply downsampled matrices or technologies like Pore-C, that the failure becomes severe. Nonzero-percentile clipping resolves this by anchoring the threshold to the observed contact distribution regardless of sparsity, making the preprocessing robust across technologies and sequencing depths.

### Whole-matrix clipping affects pixelwise but not structural metrics

Whole-matrix clipping selectively degrades pixelwise similarity metrics while leaving higher-order structural features intact. Pixelwise metrics as PCC and SSIM are severely degraded in matched comparisons, while higher-order structural metrics including insulation score correlation, TAD boundary F1, and contact decay correlation remain essentially unchanged across preprocessing conditions. This dissociation has implications in two directions. On the one hand, it suggests that biological conclusions drawn from studies using whole-matrix clipping are likely robust, since the spatial organization of the genome as reflected in domain boundaries and compartment structure is preserved for experiments like Hi-C even when the absolute contact values are heavily distorted. The extensive literature built on Hi-C data processed with these conventions is mainly focused on a preprocessing tailored to the gold-standard Rao et al. (2014) dataset rather than to a general approach across experiment types. On the other hand, this same robustness means that structural metrics are relatively insensitive instruments for benchmarking enhancement models. A model that recovers insulation structure will score well on these metrics regardless of how well it reconstructs absolute contact frequencies. PCC and SSIM, despite their sensitivity to clipping artefacts, capture a dimension of reconstruction quality that structural metrics do not. This motivates the multi-metric evaluation framework we employ, where pixelwise, structural, and decay metrics together provide a more complete picture than any single measure. This dissociation highlights that structural robustness alone is insufficient to guarantee quantitative correctness.

We also note that SSIM requires particular care when applied to contact matrices with non-standard dynamic ranges. Specifically, the dynamic range parameter L should be computed empirically from the data being compared as the observed value range of the two matrices and not be kept fixed as in the original (Wang et al., 2004) definition. The stability constants in the SSIM formula scale with the observed data range, so severe compression of the dynamic range by whole-matrix clipping artificially inflates the sensitivity of SSIM to residual differences. SSIM comparisons are only meaningful when both matrices share the same dynamic range assumptions and reported SSIM values are not comparable across studies that differ in preprocessing choice. For contact decay reconstruction specifically, we find that Pearson correlation saturates near unity even for substantially degraded matrices due to the monotonic nature of P(s) curves and recommend Lin’s concordance correlation coefficient (DecayCCC) as a more informative alternative that jointly captures both correlation and scale agreement.

### Normalization at inference time requires consistent use of input-derived statistics

A subtle but consequential source of data leakage arises from how the low-resolution input is normalized at inference time. During training, the high-resolution target matrix is available, and its statistics as the maximum value and percentile thresholds can be used to define the normalization factor per chromosome. However, at inference time only the low-resolution input is available. If the model was trained with normalization parameters derived from the high-resolution matrix, those parameters constitute a form of data leakage. Such inconsistencies compromise the interpretability of reconstruction results and can artificially inflate model performance. They encode information about the target that would not be accessible in a real use case. If instead a fixed constant is used, the model implicitly learns a scaling relationship between LR and HR rather than a true structural reconstruction, since the LR input is not compressed into the same dynamic range as the HR target. WE propose normalizing the LR input using statistics derived from the LR matrix itself, applied consistently between training and inference. In practice, this means that clipping thresholds and scaling factors must be computed from the low-resolution input alone and not from fixed constants such as 255 and especially not from the high-resolution target matrix.

Precise normalization behavior at inference time varies across published implementations. In CCUT, we normalize each matrix using per-chromosome statistics derived from that matrix alone, applied consistently during both training and inference. We believe that this is one possible approach, but we note that a rigorous empirical evaluation of the downstream effects of different inference-time normalization strategies and their interaction with model generalization across sequencing depths remains an open problem that the field would benefit from addressing systematically.

### CCUT as a framework for Pore-C enhancement and preprocessing standardization

#### CCUT serves as a proof-of-principle that physically consistent preprocessing enables meaningful reconstruction of chromatin architecture from sparse data

By providing a unified framework with a fixed, statistically appropriate preprocessing pipeline and a comprehensive evaluation suite, CCUT enables fair comparison of different architectures under identical conditions. Crucially, existing architectures that have demonstrated strong performance on Hi-C data can be incorporated into the CCUT framework and trained under the corrected preprocessing.

HiCNet, as our primary benchmark implementation within CCUT, demonstrates that the framework can recover chromatin structure from deeply downsampled Pore-C data. The consistent improvement across pixelwise, insulation, and decay metrics across chromosomes with domain organization suggests generalization beyond the specific training conditions, though systematic evaluation across a wider range of genomic contexts and experimental conditions remains necessary. The non-linear relationship between downsampling ratio and reconstruction quality, where substantial coverage reduction produces only modest metric degradation, suggests that the inherent structure of chromatin interaction data provides strong inductive constraints that a well-designed model can exploit.

### Limitations and future directions

Some limitations of the current work should be acknowledged. HiCNet is optimized for the sequencing depths and experimental conditions represented in the training data; generalization to substantially different protocols or organisms may require retraining or fine-tuning, though the relatively low computational cost of the model makes this feasible on consumer-grade hardware. Genomic regions with extreme GC content or complex repetitive elements present challenges for both sequencing and computational reconstruction that are not specific to our approach but are worth noting as areas where performance may be reduced. The absence of existing Pore-C enhancement benchmarks means that our evaluation is self-contained rather than comparative; as the field develops, direct comparisons with future methods will be important for understanding the relative strengths of different approaches.

Looking forward, CCUT’s modular design accommodates the integration of new and existing architectures, loss functions, and data types as the field evolves. The simultaneous capture of methylation information by Pore-C opens the possibility of incorporating epigenetic signals as auxiliary inputs or targets, which could further improve reconstruction quality given the known relationship between chromatin organization and DNA methylation. Extension to single-cell data, where sparsity is even more extreme and the preprocessing challenges we characterize here are correspondingly more severe, represents a particularly promising direction. The standardized preprocessing and evaluation pipeline provided by CCUT offers a common reference for these developments, directly addressing the reproducibility challenges that currently impede objective comparison across the field.

Since CCUT retains information associated with short range genomic contacts, i.e. at the level of TADs and beyond, it represents an excellent tool for studying the impact of (changes in) local physical properties of chromatin on its 3D organization by comparison of upsampled matrices to maps generated by KMC models or molecular dynamics simulations. In this work, the modeling treats the case for which the persistence length (as a measure for the chromatin stiffness) is small compared to the length of a chromatin segment consistent with a single bin. As explained above, the value of the persistence length determines the scale at which the model operates: by taking larger values, we automatically consider smaller length scales, at which the chromatin stiffness contributes to determining extrusion dynamics and contact probabilities. We note that by including additional reaction pathways, the KMC model is readily extended to account for variation in stiffness with genomic position, as well as with processes that likely affect the polymer properties, such as methylation and histone modification. More broadly, this framework opens the possibility of systematically linking chromatin contact data to quantitative polymer models across experimental conditions.

## Materials and Methods

### Data and Experimental Design

Pore-C contact frequency matrices are produced from experiments in which a large population of cells is simultaneously processed, capturing the three-dimensional spatial proximity of genomic loci across the nucleus. The result is a symmetric matrix *H_nxn_* where each entry H(i,j) represents the observed contact frequency between genomic bins i and j at a given resolution, with higher values indicating greater spatial proximity in the three-dimensional nuclear environment. Because the matrix reflects contacts averaged across all sequenced cells, it encodes the population-level chromatin conformation at the resolution determined by sequencing depth and bin size.

For this study we used Pore-C data from Deshpande et al. (2022), obtained from GEO accession GSM4490691 and processed with the Oxford Nanopore best practices Nextflow pipeline wf-Pore-C to pairs format. Contact matrices were generated from the pairs files using the cooler library (Abdennur & Mirny, 2020) at a base resolution of 10,000 bp and coarsened into multi-resolution cooler files. Raw integer counts were retained without balancing for all downstream processing. The data were split by chromosome into a training set (chromosomes 1-15), a validation set (chromosomes 16-19) and a held-out test set (chromosomes 20-22), with chromosomes 16-19 also serving as the validation set during hyperparameter optimization.

### Downsampling

To simulate reduced sequencing depth in a statistically principled way, we generated low-resolution training inputs by binomial downsampling of the raw count matrix. For each bin (i,j) with observed count c, a downsampled count c′ is drawn as:

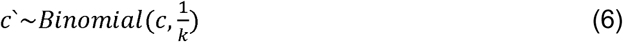

where k is the downsampling factor. This is statistically correct because each sequenced read pair is an independent Bernoulli trial. Sequencing at 1/k depth means each read is retained with probability 1/k, and the resulting bin count follows a Binomial distribution with these parameters. This per-bin Binomial approach is more accurate than uniform random sampling across the full upper triangle, which treats the matrix as a flat bag of reads without regard to the independence of different loci. For sparse bins with counts of 1-3, the distinction is practically meaningful as for high-count bins the two approaches converge. All training used a downsampling factor of k=16. Downsampled matrices were generated on-the-fly during training from the single high-resolution cooler file, either as a fixed pre-computed realization per chromosome (deterministic mode, used for evaluation) or as a fresh Binomial draw per training step (stochastic mode, used for data augmentation during training).

### Normalization Pipeline and Invertibility

Raw contact count matrices exhibit heavy-tailed distributions driven by a small number of very high-frequency near-diagonal entries. Applying a log transform directly to unclipped counts does not resolve this: extreme values are compressed but not removed, and minor differences in log space correspond to large differences in count space upon inversion, which destabilizes reconstruction. A clipping step prior to the log transform is therefore necessary.

For each chromosome, we compute the 99.95th percentile of nonzero entries only, treating zero-valued bins as structural absences rather than observations. The full matrix is then clipped at this threshold. This nonzero-percentile strategy ensures that the clip threshold reflects the actual observed contact distribution rather than the zero mass, which would otherwise dominate the percentile calculation and collapse the threshold into the bulk of the nonzero signal. By construction, clipping at the 99.95th nonzero percentile retains 99.95% of observed contact information exactly, with only the top 0.05% of nonzero entries truncated to the threshold value; we therefore characterize this step as near-lossless.

Following clipping, we apply a log1p transform, to the clipped matrix. This compresses the dynamic range of the nonzero signal into a well-conditioned range suitable for neural network training while mapping zero entries exactly to zero. Each chromosome is then scaled by its per-chromosome maximum of the log1p-transformed matrix, producing values in [0,1]. Because the minimum of the log1p-transformed clipped matrix is always zero, this reduces to a division by the per-chromosome maximum; no minimum subtraction is required. This pipeline admits a well-defined inverse. Given a model prediction in [0,1], the inverse transform proceeds as: multiply by the stored per-chromosome maximum, apply expm1, i.e. exp(x)−1, and clip any negative values to zero to handle floating-point deviations. The result is a reconstructed matrix in the original count space, accurate up to the 99.95th nonzero percentile threshold. This invertibility means that model predictions can be directly compared to and interpreted in terms of the original experimental count data.

The per-chromosome maximum used for scaling is computed from the high-resolution matrix during training. At inference time, where no high-resolution matrix is available, the per-chromosome maximum is computed from the low-resolution input matrix and used consistently for both forward and inverse transforms, ensuring that normalization does not require any statistics derived from the reconstruction target.

### Evaluation Metrics

For chromatin contact matrix <u>reconstruction,</u> we employ a combination of traditional image quality metrics and domain-specific structural metrics. Given a reference matrix H and a reconstructed matrix K of dimension m×n, the pixelwise metrics are defined as follows:

#### Mean Absolute Error (MAE) / L1 Loss

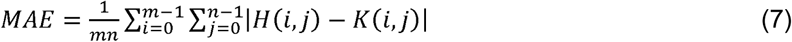

#### Mean Squared Error (MSE) / L2 Loss

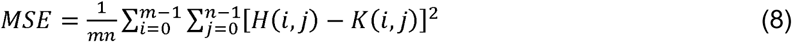

#### Peak Signal-to-Noise Ratio (PSNR)

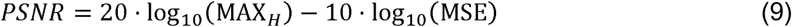

where MAX_H_ is the maximum pixel value of the image.

#### Structural Similarity Index measure (SSIM)

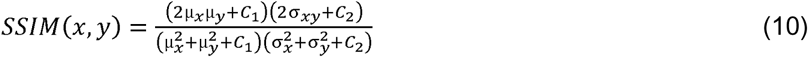

where μ*_x_*, μ*_y_* are means, 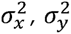 are variances, σ*_xy_* is covariance, and *C_1_ = (K_1_L)^2^*, *C_2_ = (K_2_L)^2^* are constants with *K_1_ = 0.01*, *K_2_ = 0.03*, and *L* being the dynamic range of pixel values.

These metrics capture complementary aspects of reconstruction quality: MAE and MSE are sensitive to absolute differences and therefore reflect recovery of high-intensity near-diagonal contacts including chromatin loops; SSIM captures local structural similarity and serves as a proxy for domain and compartment organization; and PSNR summarizes overall signal quality relative to the noise floor.

In addition to pixelwise metrics, we evaluate three domain-specific structural properties. Insulation score correlation measures the Pearson correlation between insulation profiles computed from the reference and reconstructed matrices, with insulation scores reflecting the degree of local chromatin compaction at each genomic position (Crane et al., 2015). TAD boundary F1 measures the overlap between boundary sets called from the reference and reconstructed insulation profiles at a fixed threshold, providing a discrete measure of domain structure recovery. Contact decay correlation measures the agreement between the P(s) curves (genome-wide mean contact frequency as a function of genomic distance) of the reference and reconstructed matrices.

For contact decay specifically, we find that Pearson correlation saturates near unity even for substantially degraded matrices due to the monotonic nature of P(s) curves and does not discriminate between poor and good reconstructions. We therefore use Lin’s concordance correlation coefficient (CCC) (Lin, 1989) as the primary P(s) metric, as it jointly measures both correlation and agreement in scale and location and is sensitive to amplitude mismatches that Pearson correlation ignores.

### HiCNet Architecture

We constructed HiCNet, a hierarchical generative adversarial network adapted for chromatin contact matrix enhancement, based on the HiNet architecture (Chen et al., 2021). The generator consists of two stages with identical U-shaped architectures, each processing the input through multiple levels of downsampling and upsampling operations with skip connections and dense blocks at each hierarchical level to preserve both local and global features. Supervised Attention Modules (SAM) are incorporated between stages to guide feature selection. Matrix symmetry, which is a structural requirement of contact data, is enforced through a weighted symmetrization of the output.

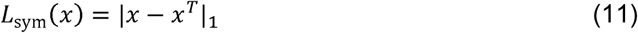

The discriminator employs a series of convolutional layers with instance normalization and LeakyReLU activations, culminating in an adaptive average pooling layer for final classification.

### Hyperparameter Optimisation

Hyperparameters for HiCNet were optimized using Optuna (Akiba et al., 2019) with 30 trials. The search space covered architecture parameters (feature width and network depth), generator and discriminator learning rates, loss component weights for pixel loss, structure consistency loss, insulation loss, and distance decay loss, loss function type (L1, MSE, or Huber), GAN configuration (enabled or disabled, gradient penalty weight, label smoothing coefficient), batch size, and early stopping patience. Within each trial, training was terminated by early stopping monitored on validation SSIM. The trial achieving the highest validation SSIM at the point of early stopping was selected as the final model configuration. **Training Protocol**

#### Structure Consistency Loss

To preserve local chromatin interaction patterns, we introduce a structure consistency loss that operates on local neighborhoods:

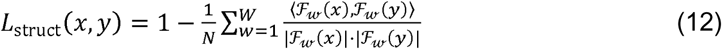

where *F_w_* extracts features from a window *w*, *W* is the total number of windows, and ⟨·, ·⟩ denotes the inner product.

#### Adversarial Loss

For our GAN-based approach, we employ a Wasserstein loss with gradient penalty:

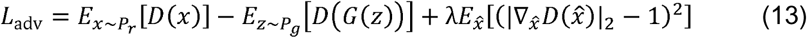

where *D* and *G* are the discriminator and generator networks respectively, *P_r_* is the real data distribution, *P_g_*is the generated data distribution, and *x*☐ represents interpolated samples between real and generated data.

Training used the Adam optimizer with generator and discriminator learning rates, gradient penalty weight, and label smoothing determined by the Optuna search. To ensure training stability, label smoothing was applied to discriminator targets to prevent overconfidence and reduce adversarial oscillations, and gradient clipping was applied with a maximum norm of 1.0 to both networks to prevent gradient explosion. Early stopping was monitored on validation SSIM with patience determined during hyperparameter search. Each session ran for a maximum of 50 epochs. The final combined loss used during training was:

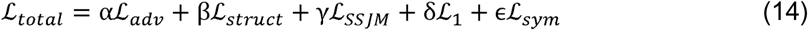

where the high weighting of the pixelwise L1 loss ensures structural fidelity and the structure consistency loss weighting preserves local chromatin interaction patterns. The model was implemented in PyTorch and trained on NVIDIA V100 GPUs using 6 parallel data loading workers.

### Cell Culture

The human lymphoblastoid cell line HG002 (Epstein–Barr virus–transformed B lymphocytes) was cultured in suspension in RPMI-1640 medium (Thermo Fisher) supplemented with 2 mM L-glutamine (GlutaMAX-I”, Gibco) and 10–15% heat-inactivated fetal bovine serum (FBS, Thermo Fisher). Cells were maintained at 37 °C in a humidified incubator with 5% CO₂. Cultures were maintained at densities between 2 × 10D and 1 × 10D cells mL⁻¹ and passaged every 2–3 days. Cell density and viability were determined by trypan blue exclusion using a hemocytometer or automated cell counter (Thermo Fisher), and only cultures with viability >85% were used for experiments. For routine passaging, cells were reseeded at approximately 2.5 × 10D cells mL⁻¹ in fresh pre-warmed complete medium.

### *C. elegans* preparation for Pore-C

Wild type *C. elegans* (N2 (Bristol)) was grown at 20°C on NGM high peptone agar plates with OP50. Embryos from 50.000 gravid adults were isolated by bleaching and maintained in H2O for 10 minutes. The sample was dounced 80X in a stainless-steel tissue grinder (Wheaton™ Dounce Dura-Grind™ Tissue Grinder, DWK Life Sciences), then centrifuged at 100g for 1 minute. Supernatant with cells was saved and the pellet resuspended in 1ml of H2O and dounced for 40 more times. Debris was removed by centrifugation at 100g for 1 minute and the supernatant with the cells from both rounds of homogenization were pooled together. Cells were then centrifuged (500g for 5 minutes) and washed 2 times with PBS. Cells were fixed at RT for 10 minutes with 1% formaldehyde solution in PBS. Formaldehyde was quenched by incubation with glycine 125 mM for 5 minutes at RT, and 10 minutes on ice. Fixed cells were washed 2 times with PBS at 4°C in a 15ml tube (5 minutes at 500g) and once in a 2ml tube. After a last centrifugation of 5 minutes at 500g the supernatant was removed and the cell pellet was snap frozen in liquid nitrogen.

### Pore-C

Pore-C library preparation was performed following the restriction enzyme Pore-C protocol from Oxford Nanopore Technologies with minor modifications (https://nanoporetech.com/document/extraction-method/restriction-pore-c). Briefly, approximately 5 × 10D cells were crosslinked with 1% formaldehyde and the reaction was quenched with glycine. Cells were permeabilized and chromatin was denatured prior to in situ restriction digestion using NlaIII. Digestion was carried out at 37 °C for 18 h to fragment chromatin while preserving spatial proximity of interacting loci. Following digestion, restriction enzymes were inactivated and proximity ligation was performed using T4 DNA ligase at 16 °C to generate concatemers representing higher-order chromatin interactions. Proteins were degraded by Proteinase K treatment and crosslinks were reversed. DNA was subsequently purified by phenol–chloroform extraction followed by ethanol precipitation, resuspended in TE buffer, and quantified using the Qubit dsDNA HS assay. Sequencing libraries were prepared using the Oxford Nanopore Pore-C kit (SQK-LSK114) according to the manufacturer’s instructions and sequenced on an Oxford Nanopore platform.

## Availability of code and data

All scripts, except for the KMC model, and the tool can be accessed via GitHub (https://github.com/stasys-hub/CCUT). The Pore-C data generated in this study (HG002 and *C. elegans*) have been deposited in NCBI GEO under accession number GSE326445 (https://www.ncbi.nlm.nih.gov/geo/query/acc.cgi?acc=GSE326445). For this study, we used Pore-C data from Deshpande et al. (2022), which was obtained from GEO GSM4490691 (https://www.ncbi.nlm.nih.gov/geo/query/acc.cgi?acc=GSM4490691) and processed with the Oxford Nanopore best practices Nextflow pipeline: wf-Pore-C (https://github.com/epi2me-labs/wf-Pore-C) to pairs format. The Micro-C Dataset from was obtained from the 4dn Portal with the accession number: 4DNFI1O6IL1Q. The script of the KMC model is available upon reasonable request.

## Authors’ Contributions

S.S. wrote the manuscript, developed methodology, and performed computational analysis. M.M. contributed to data processing, data analysis and writing the manuscript. A.S. generated the Pore-C and Hi-C data, R.H.R. generated the *C. elegans* data. S.W. contributed to the deep learning implementation. M.W. provided deep learning expertise. S.G. conceived the study and supervised the work. S.Sch. contributed to interpreting the data and editing the manuscript. J.J.M. conceived the KMC model and performed the simulations. J.P. supervised the *C. elegans* related work and interpretation.

## Acknowledgements

SG, SS, JP and JJM acknowledge funding by SFB 1551 Project No. 464588647 of the Deutsche Forschungsgemeinschaft (DFG). KES acknowledges funding from the German Research Foundation (DFG) Project No. 320163632. This work was also supported by the Deutsche Forschungsgemeinschaft (DFG, German Research Foundation) through the Collaborative Research Centre (SFB) 1361 (Project-ID 393547839). SG and JP acknowledge funding from the Mainz Institute of Multiscale Modeling.

